# Comparison of drug inhibitory effects (IC_50_) in monolayer and spheroid cultures

**DOI:** 10.1101/2020.05.05.079285

**Authors:** Catherine Berrouet, Naika Dorilas, Katarzyna A. Rejniak, Necibe Tuncer

## Abstract

Traditionally, the monolayer (two-dimensional) cell cultures are used for initial evaluation of the effectiveness of anticancer drugs. In particular, these experiments provide the IC_50_ curves that determine drug concentration that can inhibit growth of a tumor colony by half when compared to the cells grown with no exposure to the drug. Low IC_50_ value means that the drug is effective at low concentrations, and thus will show lower systemic toxicity when administered to the patient. However, in these experiments cells are grown in a monolayer, all well exposed to the drug, while *in vivo* tumors expand as three-dimensional multicellular masses, where inner cells have a limited access to the drug. Therefore, we performed computational studies to compare the IC_50_ curves for cells grown as a two-dimensional monolayer and a cross section through a three-dimensional spheroid. Our results identified conditions (drug diffusivity, drug action mechanisms and cell proliferation capabilities) under which these IC_50_ curves differ significantly. This will help experimentalists to better determine drug dosage for future *in vivo* experiments and clinical trials.

## 1 Introduction

In general, drug-dose response curves are used to measure and analyze the relationship between a drug’s inhibitory capabilities associated with its respective concentrations. Inhibitory concentration curves denoted IC_*x*_ are dose-response curves that allow for determining the drug concentration required to reduce a population of viable cells by x%, when compared to the cells grown with no exposure to the drug [1]. This change in cell population size could be a result of increased cell death or suppressed cell proliferation. Drug discovery and pharmacology studies use the IC_50_ values to determine drug effectiveness (potency). Low IC-, value means that the drug is potent at low concentrations, and thus will show lower systemic toxicity when administered to the patient. The drug dose-response curves are also used to identify synergistic combination therapies and drug interactions mechanisms [2, 3, 4].

However, the typical experiments to determine the IC_50_ curves are performed in two-dimensional (2D) monolayer cell cultures in Petri dishes (Fig. 1A). The cells are covered with a medium of uniform drug concentration and grown for 72 hours which is a timeframe long enough for cells to divide 1-2 times and to observe drug effects without cells reaching confluence [5, 6]. In contrast, *in vivo* tumors develop as three-dimensional (3D) masses of tightly packed tumor cells, and thus their response to therapeutic interventions may be different than in 2D experiments. To test drug potency in *vitro* in a way to preserve geometry of typical in vivo tumors, the 3D cultures of multicellular spheroids were developed [7, 8, 9]. In these experiments, the 3D spheroids are first formed either by proliferation from single seeded cells or by aggregation of individual cells seeded together [9, 10]. The spheroids are then covered with a medium with a uniformly dissolved drug (Fig. 1B), similarly like this is done in the 2D experiments. However, a significantly different culture geometry results in a limited access to the drug inside the spheroid. There is no standard experimental procedure for assessing drug response in these 3D cultures, therefore, we designed computational models as *in silico* analogues of the 2D monolayer and a cross section through the 3D multicellular spheroid cell cultures (Fig. 1) to compare side-by-side how identical cells respond to identical drugs in these two settings starting with the same number of cells. To our knowledge, this is the first comprehensive comparison between drug efficacy simulated using analogues of 2D and 3D cell cultures.

**Figure 1:**
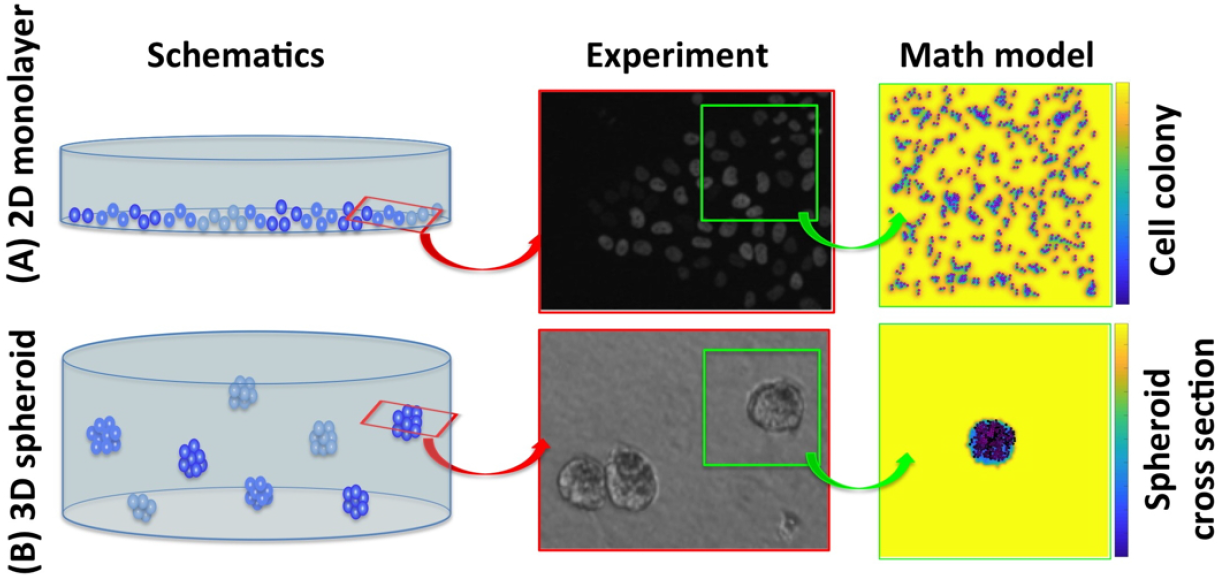
Schematics of 2D and 3D cell cultures and their mathematical analogues. **A.** A schematics of the 2D monolayer cell culture, a corresponding fluorescent microscopy image of a part of the Petri dish, and a snapshot from our *in silico* analogue model of the 2D cell culture. **B.** A schematics of the 3D multicellular spheroid culture, a corresponding bright field microscopy image of a part of the well plate, and a snapshot from our in *silico* analogue model of the cross section through the 3D spheroid. The background color represents drug concentration to illustrate local drug gradients from high concentration (yellow) to low concentration (blue). Experimental images courtesy of the Moffitt Analytic Microscopy Core.

In this paper, we first describe a mathematical framework used to model the 2D monolayer cell culture and the cross section through a 3D multicellular spheroid culture (Section 2). Numerical implementation of these models is described in Appendix 4. These two models are then used to test conditions (drug diffusivity, drug action mechanisms and cell proliferation capabilities) under which the IC_50_ values are either similar or significantly different between 2D and 3D cultures. The method of fitting the IC_50_ curves to simulated data is presented in Section 3. The analysis of results for cytotoxic drugs is presented in Section 5.1, and for anti-mitotic drugs in Section 5.2. We summarize our results with the Discussion Section 6.

## 2 The Mathematical Model

From a mathematical modeling perspective of tumor cell populations, the models can be classified into two types: continuous or discrete. Continuous models treat the population of tumor cells as a continuous density distribution, which usually is described by a system of ordinary or partial differential equations. On the other hand, in discrete models each cell in the population is represented by a discrete object (an agent) that follows a set of prescribed rules. Thus, these models are often referred as agent-based models (ABMs). Depending on the research question, the one or the other mathematical modeling approach is considered; though both have their strengths and weaknesses. For a review on the comparison of the two types of models, see [11]. A continuous model is relatively easier to analyze analytically and computationally but it fails to capture the individual cell-to-cell interactions and cellular heterogeneity. On the other hand, the ABM models focus on interactions between the individual cells but are not easy to analyze analytically, if not impossible, and the computation time increases significantly with the number of cells involved. Recently, researchers have been focusing on developing hybrid discrete-continuous models which combine both approaches with the goal of maximizing their advantages and minimizing their drawbacks. More information on the hybrid discrete-continuous models can be found in [12].

Here, we use a hybrid discrete-continuous model in which tumor cells interact physically with one another and react to a drug dissolved in a surrounding medium. The cells are modeled as individual off-lattice agents, and drug concentration is described by the continuous partial differential equation. We previously used a similar mathematical framework to model *in vivo* tumors and the emergence of drug-induced resistance [13, 14, 15, 16, 17]. Here, we adjusted this framework to model *in vitro* cell cultures, both the 2D monolayer and the cross section through the 3D spheroid. In addition to different culture geometry, we also extended the previous work by considering different mechanisms of drug action, i.e., the cytotoxic and anti-mitotic drugs. Below, we provide all model equations and spatial setups of all performed computational experiments.

### 2.1 Discrete description of tumor cells

Each cell in our model is treated as a separate entity characterized by several individually regulated properties. The position of *k^th^* cell is denoted by **X**_*k*_ (*t*), current cell age by 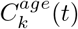, cell maturation age, which is the age when the cell is ready to divide, by 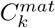, the number of nearby neighboring cells, 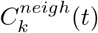, and a level of the drug accumulated inside the cell by 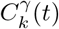. For simplicity, we assume that all cells have the same diameter *R_D_*. The state of the *k^th^* cell at time *t* is denoted by 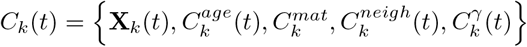. The initial state of the *k^th^* cell is 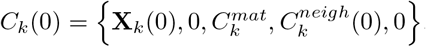.

Note that the cell maturation age, 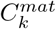 is the only cell feature that does not depend on time *t*, and it is assigned explicitly upon cell birth. However, to avoid cell synchronized divisions, we impose up to 15% of differences in the duration of the cell cycle [18]. Thus, we set

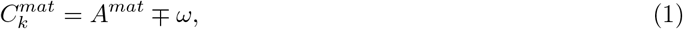

where *A^mat^* is the average maturation age for the whole cell population, and *ω* is randomly chosen from a uniform distribution from [0, 2.5] hours.

The cell *C_i_* is considered to be a neighbor of cell *C_k_* at time *t*, if it is located within the neighborhood radius *R_neigh_* from cell *C_k_*, that is:

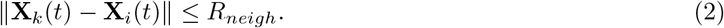

We consider here the neighborhood radius equal to two cell diameters (*R_neigh_* = 2*R_D_*) which accounts for two layers around the host cell (as in other models [14, 19]). The initial number of neighboring cells, 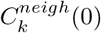 depends on the initial cell configuration (see section 2.3). As the cells divide, die or move around, the number of neighboring cells, 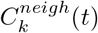, varies with time.

The cell spatial dynamics is modeled by Newton’s second law of motion:

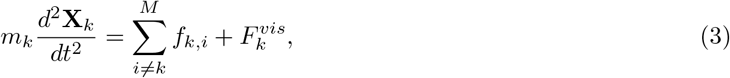

where *f_k,i_* is the interaction force between two neighboring cells *C_k_* and 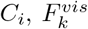 denotes the force against viscosity of the surrounding medium, and *m_k_* is the mass of the *k^th^* cell. Newton’s second law of motion has been used to address the dynamics of cell-to-cell interactions, see the review in [20] and the reference therein.

The cell interaction forces arise when two neighboring cells, *C_k_* and *C_i_*, come into too close contact. If the distance between the cells’ centers is smaller than the cell diameter *R_D_*, the repulsive Hooke’s forces *f_k,i_* and *f_i,k_* = − *f_k,i_* are exerted to preserve cells’ volumes:

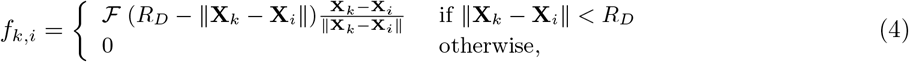

where 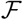 is the constant spring stiffness, and the spring resting length is equal to cell diameter *R_D_*. Since the cell can be exposed to interactions with multiple neighbors, the total force acting on the *k^th^* cell is the sum of all repulsive forces *f*_*k*,1_ + *f*_*k*,2_ + … + *f_k,M_* between the *k^th^* cell and its *M* neighbors.

Cell dynamics is thus governed by the equations of motion where the connecting springs are overdamped and system returns to equilibrium without oscillations. Hence

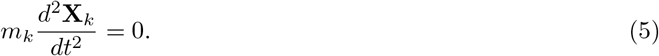

The viscous force, 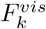 of the *k^th^* cell is modeled as proportional to its velocity. That is

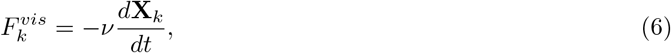

where *ν* denotes the media viscosity coefficient. Then substituting (5) and (6) into (3), we obtain that the cell motion is determined by

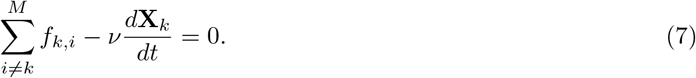

Clearly, cell’s age progresses at the same rate as time progresses, hence

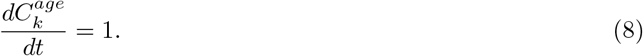

This equation has an exact solution 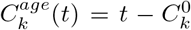 where 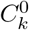 is the time at which the cell was born. However, we will keep the differential equation form for consistency with other equations in numerical implementation described in Appendix 4.

When the *k^th^* cell reaches its maturation age, 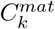, it will divide unless it is overcrowded. The overcrowding means that the number of neighboring cells, 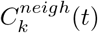, located within the neighborhood radius, *R_neigh_*, exceeds the prescribed threshold 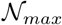. If the cell is overcrowded, its proliferation is suppressed until the space becomes available. Here, the overcrowding threshold was determined computationally to be equal to 10 cells.

Upon division of the *k^th^* cell, two daughter cells *C*_*k*_1__ (*t*) and *C*_*k*_2__ (*t*) are created instantaneously. They are placed symmetrically near the mother cell in the random direction *θ* chosen from a uniform discrete distribution [0,2π]. The locations of the daughter cells are defined by

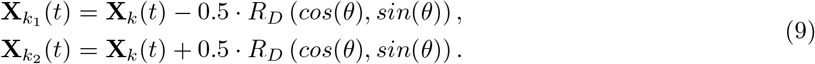

Since they are placed at the distance smaller than cell diameter, the repulsive forces between daughter cells are activated. Furthermore, this may also result in daughter cell placement near other cells. Therefore, multiple repulsive forces will be applied to resolve potential cell overlap and the cells will be pushed away until the whole cell cluster reaches an equilibrium configuration.

The age of each newly-born daughter cell is initialized to zero,

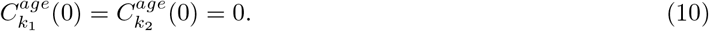

The cell maturation age is inherited from its mother cell, however, a small noise term is added to avoid synchronization of the cell cycles,

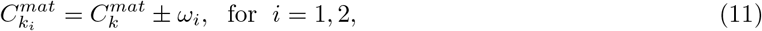

where *ω_i_* is randomly chosen from a uniform distribution [0,2.5] hours.

The level of drug accumulated by the mother cell is divided equally between the two daughter cells,

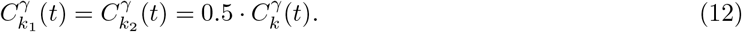

The overall rate of change in drug accumulated by the *k^th^* cell is modeled in the most simple way as absorption at a constant rate *ρ_γ_*.

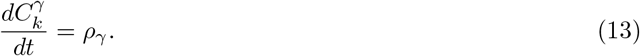

This equations has an exact solution: 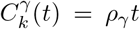, provided that drug concentrations at the locations visited by the cell are not lower that cell’s demand. Thus, special precautions will be taken in the numerical implementation described in Appendix 4 to account for these situations.

We model here two different mechanisms of drug action. In the case of a cytotoxic drug, the cell dies immediately after accumulation of a lethal dose of the drug *γ^max^*. In the case of an anti-mitotic drug, the cell dies when it attempts to divide (in the mitotic phase of the cell cycle) after accumulated drug exceeds the lethal dose *γ^max^*. The dead cells are removed from the system.

### 2.2 Continuous description of drug kinetics

The change in drug concentration *γ*(*x, t*) at location x=(*x, y*) within the domain Ω, depends on drug diffusivity and on the uptake by the tumor cells located nearby. The partial differential equation (PDE) describing the spatio-temporal drug dynamics is given by:

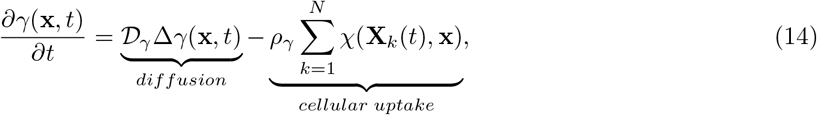

where, 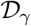 is the drug diffusion coefficient, *ρ_γ_* is the cellular uptake rate, *N* is the total number of cells located in the neighborhood of radius *R_γ_* defined by the indicator function *χ*:

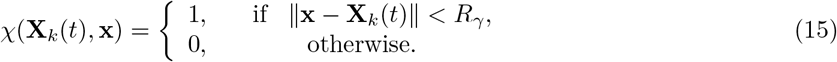

We assume that there is no loss or gain of the drug along the domain boundaries *∂*Ω. Hence, we impose Neumann-type boundary conditions

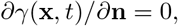

where x ∈ *∂*Ω, where **n** is the inward pointing normal.

The drug is supplied only once, at the beginning of the simulation and its concentration in the extracellular space, that is in the space outside the cells 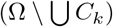, at time *t* = 0 is uniform,

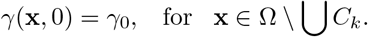

### 2.3 Initial cellular configurations

Since our goal is to compare the IC_50_ values achieved in the computational analogues of the 2D and 3D cell cultures when the same cells are exposed to the same drug, we consider two different initial configurations that correspond to these laboratory experiments. In both cases, we start with the same initial number of cells (315) that was determined computationally to ensure that the cells will not grow to confluence during the simulated 72 hours. This is consistent with laboratory experiments. We reproduced the cell monolayer culture (Fig. 2A) and the cross-section though the cell spheroid culture (Fig. 2B), respectively.

**Figure 2:**
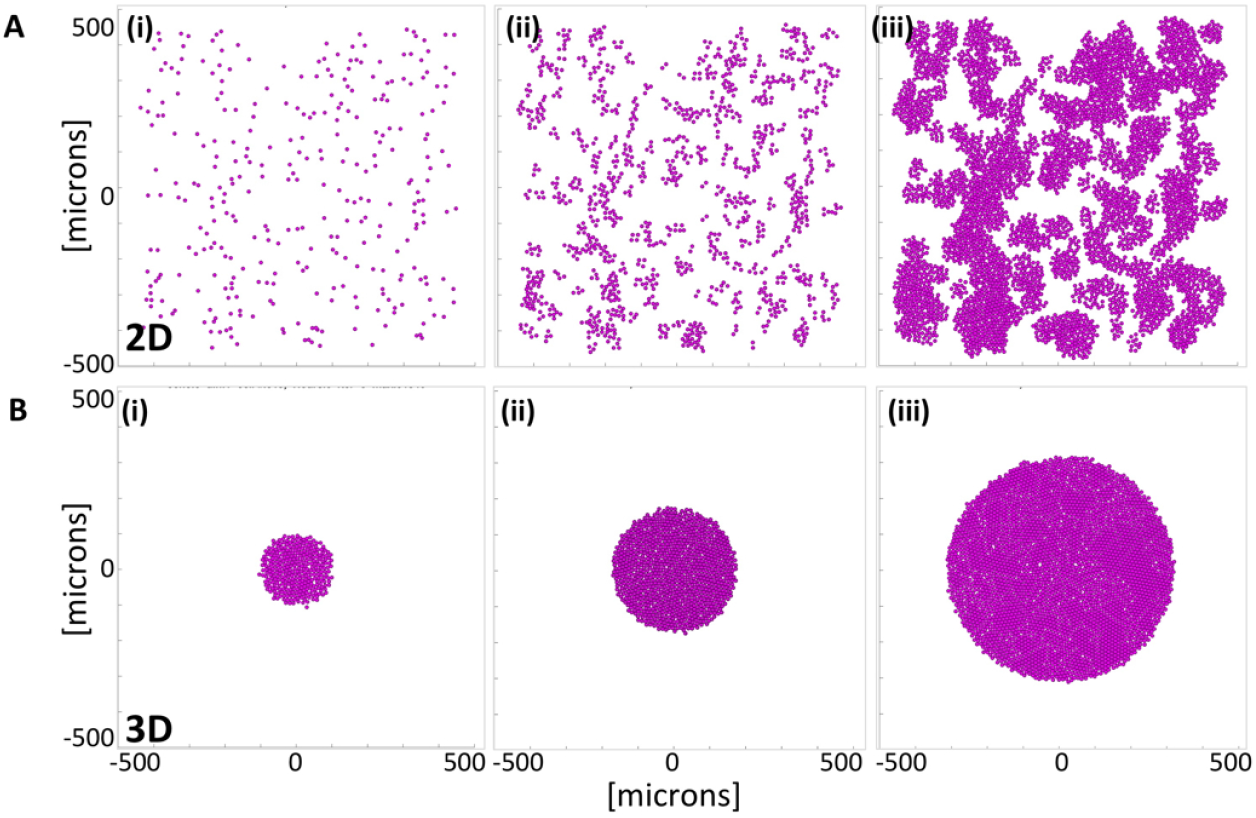
Tumor growth in 2D and 3D cell cultures with no drug (the control case). **A.** A computational model of the monolayer cell culture with sparsely seeded initial 315 cells (i), the daughter cells divide and spread throughout the domain (ii) for 72 hours of the simulated time reaching 4516 cells (iii). **B.** A computational model of the cross-section though the spheroid cell culture with initial cluster of 315 cells (i), the non-overcrowded cells divide and expand (ii) for 72 hours of the simulated time reaching 3183 cells (iii).

#### Modeling the monolayer cell culture

The two-dimensional computational model is designed to reproduce the evolution of cells seeded sparsely in the domain filled with a dissolved drug. This is an analogue of a 2D cell monolayer culture in the Petri dish where all cells are exposed to the drug dissolved in the surrounding medium [1, 2]. Each simulation starts with 315 cells that are randomly distributed within a domain. The cells are monitored for 72 hours and the final number of viable cells is recorded. During this computational experiment the cells are allowed to divide, absorb the drug and die. Upon division, the daughter cells are placed randomly nearby the mother cell and the repulsive forces are applied between overlapping cells until the new stable configuration is achieved. The cells could also become growth-arrested due to contact inhibition if their configuration reached confluence. Upon death, the cells are removed from the system which could create free space and initiate new cell divisions. Fig. 2A(i)-(iii) shows three snapshots from a monolayer simulation with no drug.

#### Modeling a cross-section of the multicellular spheroid culture

The two-dimensional computational model is designed to reproduce the central cross-sectiton of 3D multicellular spheroid culture. In these laboratory experiments, the cells are first grown in the dish until they form a packed sphere, and then are embedded into the medium mixed with a drug [21, 22]. We reproduce this experimental design by starting each simulation with 315 cells that formed a circular cell cluster. The cells are then monitored for 72 hours and are allowed to proliferate, absorb the drug or die, as in the monolayer case. However, the difference in initial configurations between these two computational models results in distinct population dynamics. In the spheroid model, the cells inside the cluster are overcrowded, and thus stop proliferating due to cellular contact inhibition. Fig.2B(i)-(iii) shows three snapshots from a multicellular spheroid simulation with no drug.

#### Drug initial distribution in the monolayer and spheroid models

The initial drug concentration *γ*(x,*t*_0_) = *γ*_0_ is uniform throughout the extracellular space outside the cells, that is in 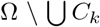. Since in the monolayer cell culture model the cells are located sparsely inside the domain, drug concentration *γ*_0_ is imposed uniformly in the whole domain (and on each grid point in the numerical implementation). In contrast, in the spheroid experiments, the drug is initially present outside the 3D culture only. To mimic the lack of the drug inside the spheroid in numerical implementation, the grid points inside the spheroid are left with no drug and drug diffusion occurs at the border of the cell cluster in the medium. However, in both computational models drug absorption by the cells leads to local depletion of the drug (see the last column in Fig.1). All physical and computational model parameters are summarized in Table 1.

**Table 1:** Physical and computational model parameters.

| **cellular & microenvironmental parameters:** |  |  |
| --- | --- | --- |
| Cell diameter | $R_D = 10\mu m$, cell radius $R = 5\mu m$ | [23, 24] |
| Spring stiffness | $\mathcal{F} = 50\mu g/\mu m \cdot s^2$ | [25, 26] |
| Mass viscosity | $\nu = 250\mu g/\mu m \cdot s$ | [27] |
| Neighborhood radius | $R_{neigh} = 2R_D = 20\mu m$ | [14, 19] |
| Overcrowding cell number | $\mathcal{N}_{max} = 10$ cells | estimated |
| Average maturation age | $\mathcal{A}^{mat} = 18, 30, 50$ hours | [6, 28] |
| **drug parameters:** |  |  |
| Diffusion coefficient | $\mathcal{D}_\gamma = 1, 10^{-2}, 10^{-4}\mu m^2/s$ | [29] |
| Drug uptake radius | $R_\gamma = 5\mu m$ | cell radius |
| Baseline drug concentration | $\bar{\gamma} = 1mM$ | |
| Initial drug concentration | $\gamma_0$ vary from 0 to $10^3mM$ | [30] |
| Cellular uptake rate | $\rho_\gamma = 5 \times 10^{-8}ng/\mu m^3s$ | $0.1\bar{\gamma}/s$ |
| Cellular death threshold | $\gamma^{max} = 2 \times 10^{-6}ng/\mu m^3$ | $2\bar{\gamma}$ |
| **numerical parameters:** |  |  |
| Domain size | $[-500, 500]\mu m \times [-500, 500]\mu m$ | |
| Mesh width | $\Delta x = \Delta y = 5\mu m$ | cell radius |
| Time step | $\Delta t = 5s$ | FTCS stability |

## 3 Fitting the Inhibitory Concentration (IC_50_) Curves

The inhibitory concentration curve (called also the drug-response curve or the IC_50_ curve) is described by the following Hill equation [31, 32]:

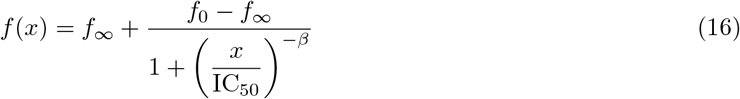

where *f*_0_ is the control effect (the plateau observed for low drug concentrations), *f*_∞_ is the background effect (the plateau observed for large drug concentrations), *β* is the curve slope, and IC_50_ value is the curve inflection point at which the drug maximal effect (*f*_0_ – *f*_∞_) decreases by 50%. The curve slope *β* is a measure of variability in drug response—the steeper the slope is the more homogeneous the drug response. The value of *f*_∞_ describes drug efficacy—the lower the *f*_∞_ the higher the beneficial effect (often denoted by *E*_max_, maximal effect). The IC_50_ value is correlated with drug potency, i.e. the amount of drug necessary to produce the effect—the lower the IC_50_ value the more potent the drug [33]. The relationship between drug potency, drug efficacy and the IC_50_ curve shape is shown in Fig.3.

**Figure 3:**
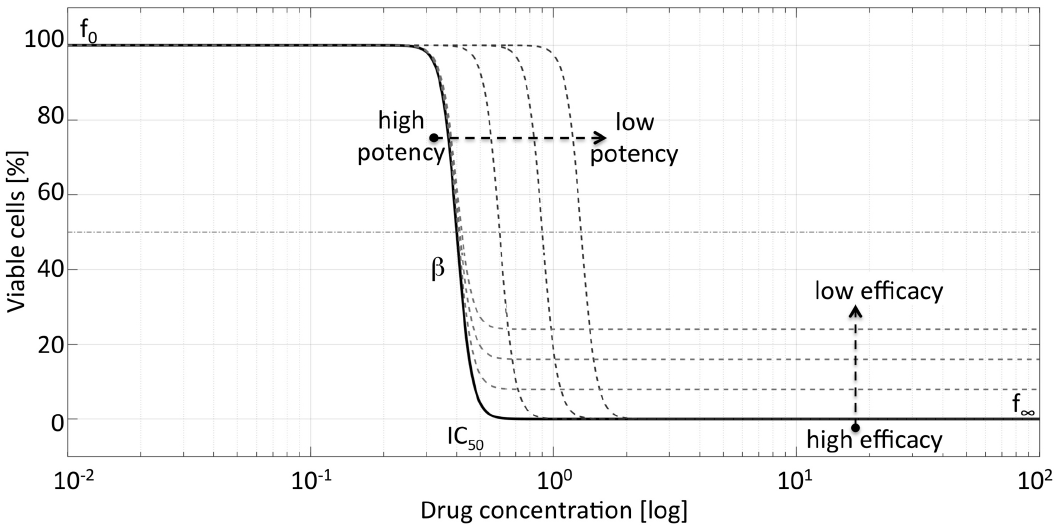
Drug potency and efficacy reflected in IC_50_ curves. The shape of the IC_50_ curve depends on the control effect (*f*_0_), variability of cell response (slope *β*), drug potency (IC_50_ value), and drug efficacy (background effect *f*_∞_).

To draw the inhibitory concentration curves and determine the half-inhibitory concentration value IC_50_, we perform computational experiments for the monolayer and spheroid cultures with initial drug concentrations varied from *γ*_0_=0 to 10^3^ mM. Each simulation starts with 315 cells arranged either in the monolayer or spheroid configuration. The total number of cells that remained after 72 hours of the simulated time is recorded. For each drug concentration, we repeat the simulation 3 times to determine the average number of viable cells. These numbers were normalized using the average count of cells grown with no drug. With this process, we obtained the normalized cell counts *f*(*γ*_0_) for each tested drug concentration *γ*_0_. For simplicity, we focus on drugs that have high efficacy, *f*_∞_ = 0, and assume that 100% of viable cells remained at end of control experiment, i.e., *f*_0_ = 100. Thus, we consider the following simplified Hill function:

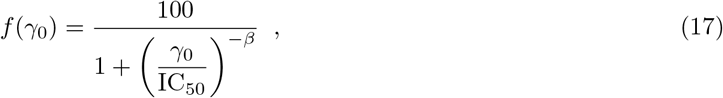

where *γ*_0_ is the drug concentration and *f* (*γ*_0_) is the normalized cell count for drug concentration *γ*_0_. The Hill coefficient *β* and the half-inhibitory concentration value IC_50_ were determined by fitting the Hill function Eq.(17) to the simulated data using the MATLAB built-in function fit that minimizes

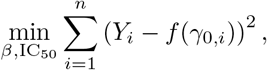

where 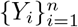 are the simulated normalized cell count data at the tested drug concentrations *γ*_0,*i*_ and 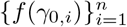 are the normalized cell counts given by the Hill function Eq.(17).

## 4 Numerical Implementation of Model Equations

Our computational model combines the off-lattice individual cells with the continuous PDE for the drug concentration. The drug concentration equation Eq.(14) is numerically solved on the regular Cartesian grid **x** = (*x,y*), while cells’ positions are defined off-lattice 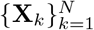. The exchange of information between these two computational structures takes advantage of the indicator function Eq.(15).

A finite difference scheme is used to approximate the solution of the differential equations modeling cell relocation and drug kinetics. Let Δt denote the time step, and let cell velocity be discretized by first order difference equation, then the cell motion Eq.(7) is discretized as

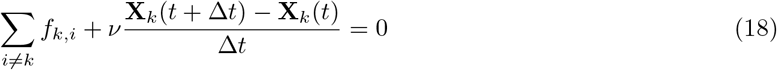

Thus, the location of the *k^th^* cell at the next time step (*t* + Δ*t*) can be determined by

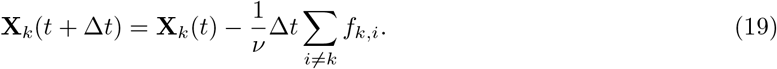

Using the same finite difference approximation for Eq.(8), the age of the *k^th^* cell at the next time step is given by

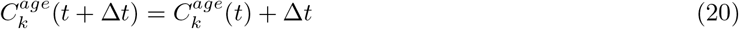

Similarly, let the rate of change of the drug accumulated in the *k^th^* cell be discretized by the first order finite difference scheme, then Eq.(13) becomes,

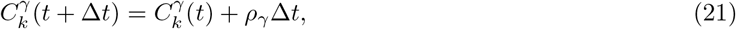

However, if there is not enough drug in cell vicinity (within the cell radius *R_D_*) to be absorbed by the *k^th^* cell, then the cell would uptake all the drug available nearby. Hence, the drug accumulated by the *k^th^* cell at time (*t* + Δ*t*) will be computed by

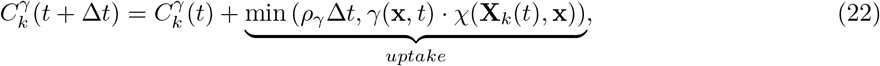

The domain Ω is discretized into a square grid with equal spacing between grid points Δ*x* = Δ*y*. Solution of the reaction-diffusion equation Eq.(14) modeling drug concentration in the domain is then approximated using a forward finite-difference approximation in time and centered finite-difference approximation in space. Thus, an approximation of the solution of Eq.(14) is obtained by the following numerical method,

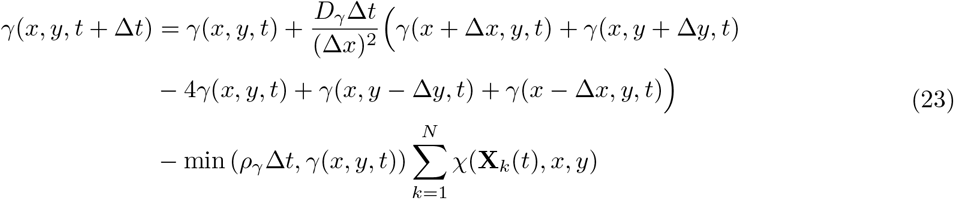

Numerical stability of this forward in time centered in space (FTCS) method is ensured by satisfying the following condition:

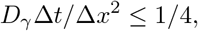

which is satisfied for the maximum diffusion value considered in this study (see Table 1 for parameter values), and the chose time step and grid size. Thus the numerical method FTCS for all parameter values is stable.

Other numerical methods could be used to approximate Eq.(14), such as an unconditionally stable fully implicit method or an IMEX method that implicit-explicit combines an implicit scheme for diffusion with an explicit scheme for the reaction terms [40, 41]. This, however, would impose higher computation costs.

In the case, when cell’s demand for drug absorption is higher than the drug available in the medium, all available drug will be depleted, thus the cellular uptake component is equal to the minimum between the drug demanded by the cells and the drug available:

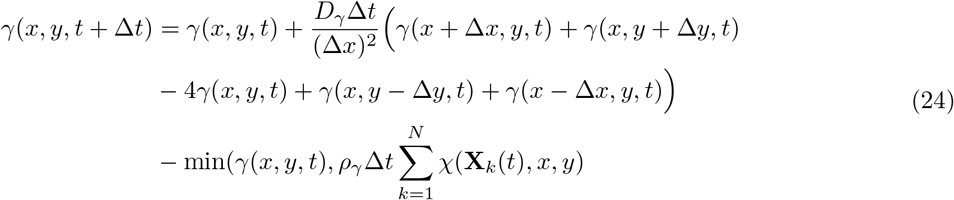

## 5 Results

Our main goal is to examine under which conditions the IC_50_ values calculated from 2D monolayer and 3D spheroid culture data are similar, and under which conditions they are distinct. This will allow us to determine, whether the typical 2D cell culture experiments are sufficient in assessing efficacy of the anticancer drugs, or if the experiments should be carried using the 3D spheroid cultures. In particular, we are interested how the IC_50_ values depend on drug mechanism of action, on drug diffusivity, and on vital properties of the tumor cells. We consider here three cell lines that differ in their doubling times and span division ages consistent with reported experimental data [6, 34]. We perform simulations for three distinct drug diffusivity values that correspond to therapeutic compounds used in the clinic, such as small molecule drugs, nanoparticles or antibodies, as reported in literature [29, 35]. We also take into consideration two distinct killing mechanisms characteristic for clinically applicable drugs [36, 37]: the cytotoxic drugs (Section 5.1) and the anti-mitotic drugs (Section 5.2). In order to cross-examine the results for both drugs, we assume that each drug is absorbed by the cells at the same constant rate and the same drug concentration is lethal for the cells.

### 5.1 Cytotoxic Anti-Cancer Drugs

In the case of cytotoxic drugs, cell survival is regulated only by the level of the absorbed drug, and the cell dies if the amount of the accumulated drug exceeds the predefined threshold. We first examined whether the IC_50_ values for the 2D and 3D cultures depend on how fast the drug is diffusing within each cell culture. Next, we examine whether the drug-induced cell death depends on how fast the cells are dividing.

#### The impact of drug diffusivity on IC_50_ values in 2D and 3D cell cultures

We considered three distinct diffusion coefficients: 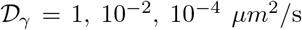. In each case, we perform several computational simulations using initial drug concentrations *γ*_0_ between 0 and 10^3^ mM for both a monolayer and a spheroid cultures. These results are summarized in Fig. 4A-C.

**Figure 4:**
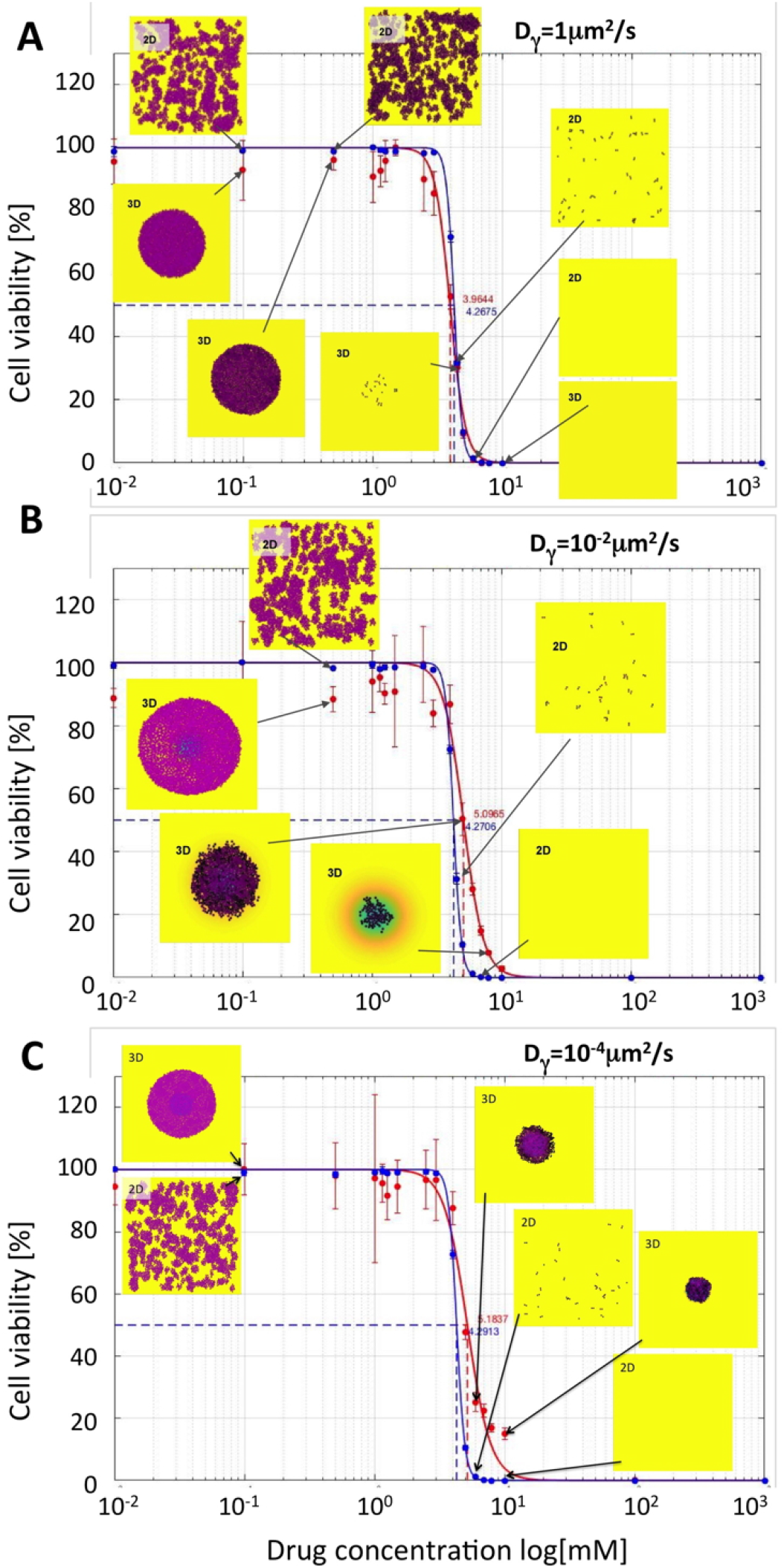
Comparison of IC_50_ curves and values for cytotoxic drugs. IC_50_ curves for monolayer cultures (blue curves) and cross-sections though spheroid cultures (red curves) and their corresponding IC_50_ values are shown for the following diffusivity values: **A.** D_*γ*_ = 1*μm*^2^/*s*; IC_50_=4.2675 for 2D and 3.9644 for 3D cultures; **B.** D_*γ*_ = 10^−2^*μm*^2^/*s*; IC_50_=4.2706 for 2D and 5.0965 for 3D cultures; **C.** D_*γ*_ = 10^−4^*μm*^2^/*s*; IC_50_=4.2913 for 2D and 5.1837 for 3D cultures. The vertical lines show standard deviation values. Insets show final cell configurations for the selected drug concentrations. Cell colors indicate the level of absorbed drug (low-pink, high-black). Background colors indicate the level of the remaining drug (high-yellow, mediumgreen, low-blue). The drug was supplied uniformly only once, at the beginning of each simulation. The cell maturation time for these simulations was 18 hours.

For each drug diffusivity, the simulations are repeated 3 times, and the average number of viable cells is used to calculate the IC_50_ curves, separately for the 2D monolayer cultures (blue curves) and 3D multicellular spheroid cultures (red curves). All IC_50_ curves have quite similar shapes and the corresponding IC_50_ values are of the same order of magnitude with less than 20% of difference. It can be observed, that with the diminishing diffusion coefficients, the IC_50_ values for the spheroid cultures are slightly higher than for the 2D cultures that is consistent with the fact that slowly diffusing drugs are not able to effectively penetrate the tightly packed cell clusters. Therefore, the cells in the 3D spheroids could survive better the drug insult. This is also confirmed by the final cellular configurations showed in the insets in Fig. 4. For higher concentrations of slowly diffusing drugs, some cells in the 3D spheroids have survived after 72 hours of exposure to the drug, while no cells remained in the 2D cell cultures exposed to the same drug concentrations. For example, when cells were exposed to the initial drug concentration of *γ*_0_ = 10 mM, there were no cell remaining in both 2D and 3D experiments for the highest diffusion case of 1*μm*^2^/s (Fig. 4A), while 105 and 315 cells remained in the 3D spheroid model for diffusion of 10^−2^*μm*^2^/s (Fig. 4B) and 10^−4^*μm*^2^/s (Fig. 4C).

In summary, slower drug diffusion resulted in more pronounced changes in the IC_50_ values for the spheroid model in comparison with the IC_50_ values for the monolayer model, which were not significantly different.

#### The impact of cell maturation ages on IC_50_ values in 2D and 3D cell cultures

Since the amount of the absorbed drug depends on how long the cell is exposed to it, we examined the relationship between the IC_50_ curves as the average cell maturation age *A^mat^* increases from 18, to 30, to 50 hours, while all other parameters were kept fixed. These results are shown in Fig. 5A-C.

**Figure 5:**
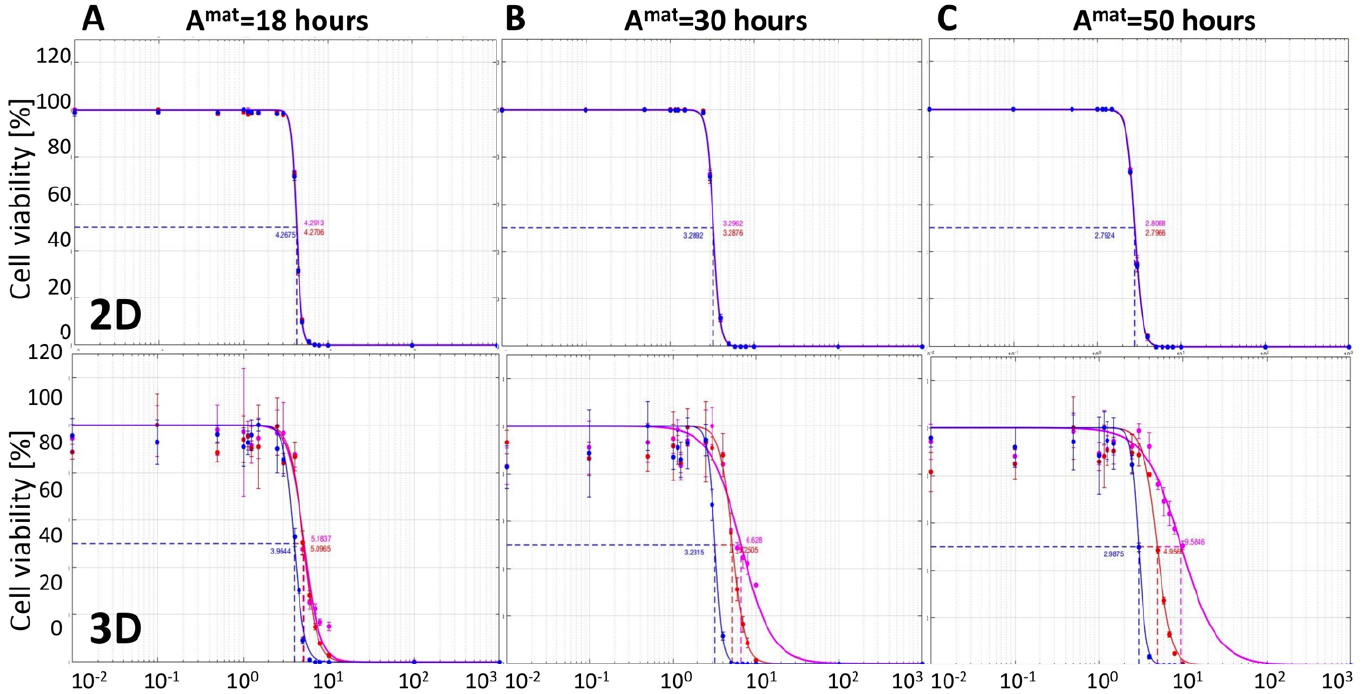
Comparison of IC_50_ curves for three cell lines of different maturation ages. IC_50_ curves and values for drugs with diffusion coefficients of D_*γ*_ = 1*μm*^2^/*s* (blue), 10^−2^*μm*^2^/*s* (red), and 10^−4^*μm*^2^/*s* (magenta) in the monolayer culture model (top) and the cross-section though the spheroid culture model (bottom) for the maturation times of **A.** *A^mat^* = 18 hours, **B.** *A^mat^* = 30 hours, and **C.** *A^mat^* = 50 hours. All data are presented after 72 hours of the simulated time.

For each maturation time, we perform simulations for both the monolayer (top row in Fig. 5) and spheroid (bottom row) cultures. We consider three diffusion coefficients: 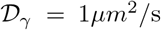 (blue curves), 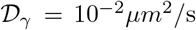 (red curves), and 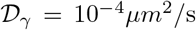 (magenta curves). The corresponding IC_50_ values for all simulated cases are summarized in Table 2. These results indicate that in the monolayer model cell maturation time and drug diffusivity do not effect the IC_50_ curves, and thus have insignificant role in inhibiting growth of the whole cell population. On the other hand, in the spheroid model, the changes in the IC_50_ values are more pronounced. For a fixed cell maturation age, the IC_50_ values differ at lease two-fold between the fastest and the slowest diffusing drug. Moreover, in the case of slowly proliferating cells, this difference is almost five-fold. For faster diffusing drugs, the IC_50_ values are similar despite the differences in cell maturation ages, but in the cases of slower spreading drugs, the corresponding IC_50_ values increase with the increased maturation.

**Table 2:** IC_50_ values for cytotoxic drug diffusion coefficients *D_γ_* and cell maturation ages *A^mat^*, for both monolayer and spheroid *in silico* cultures.

| Summary of $IC_{50}$ values in the monolayer model | | | |
| --- | --- | --- | --- |
| $D_\gamma \mu m^2/s$ | $A^{mat} = 18$ hours | $A^{mat} = 30$ hours | $A^{mat} = 50$ hours |
| 1 | 4.2675 | 3.2892 | 2.7924 |
| $10^{-2}$ | 4.2706 | 3.2876 | 2.7966 |
| $10^{-4}$ | 4.2913 | 3.2962 | 2.8088 |
| Summary of $IC_{50}$ values in the cross-section spheroid model | | | |
| $D_\gamma \mu m^2/s$ | $A^{mat} = 18$ hours | $A^{mat} = 30$ hours | $A^{mat} = 50$ hours |
| 1 | 3.9644 | 3.2315 | 2.9875 |
| $10^{-2}$ | 5.0965 | 5.2505 | 4.9565 |
| $10^{-4}$ | 5.1837 | 6.6280 | 9.5846 |

In summary, the 2D monolayer model is not able to differentiate between the inhibitory effects of cytotoxic drugs of different diffusivity for a range of tumor cells with diverse proliferation dynamics. This is because all cells in the monolayer culture are equally exposed to the drug. On the other hand, in the 3D spheroid culture model, both factors: drug diffusivity and frequency of cell proliferation affect the IC_50_ values noticeably, with values increasing for slower diffusing drugs and for slower proliferating cells. This is an effect of a more realistic tumor geometry in which tightly packed cells are more difficult to penetrate by the drug dissolved in the surrounding external medium.

### 5.2 Anti-Mitotic Anti-Cancer Drug

The anti-mitotic drugs are designed to interfere with microtubule functions which are essential in the cell division process. Cells exposed to these drugs can not progress to the mitotic phase of the cell cycle and die. Therefore, cells’ survival depends on both the level of the accumulated drug and the current cell cycle phase. In particular, if the absorbed drug exceeds the predefined threshold, the cell remains viable until it attempts to divide. For the anti-mitotic drugs, we first examined whether the IC_50_ values for the 2D and 3D cultures depend on drug diffusivity within the 2D and 3D cell cultures. Next, we tested whether the drug-induced cell death depends on cell maturation age, that is on how fast the cells are dividing.

#### The impact of drug diffusivity on IC_50_ values in 2D and 3D cell cultures

As in the previous case, we considered three different drug diffusion rates: 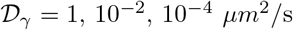, and performed several computational simulations using initial drug concentrations *γ*_0_ between 0 and 10^3^ mM. Each simulation are seeded with 315 cells arranged either sparse in the domain (a monolayer culture) of in a circular configuration (a cross section through a spheroid culture). For each combination of parameters, we perform 3 simulations and the average data is used to generate the IC_50_ curves for the 2D monolayer cultures (blue curves) and 3D multicellular spheroid cultures (red curves). These results are presented in Fig. 6A-C.

**Figure 6:**
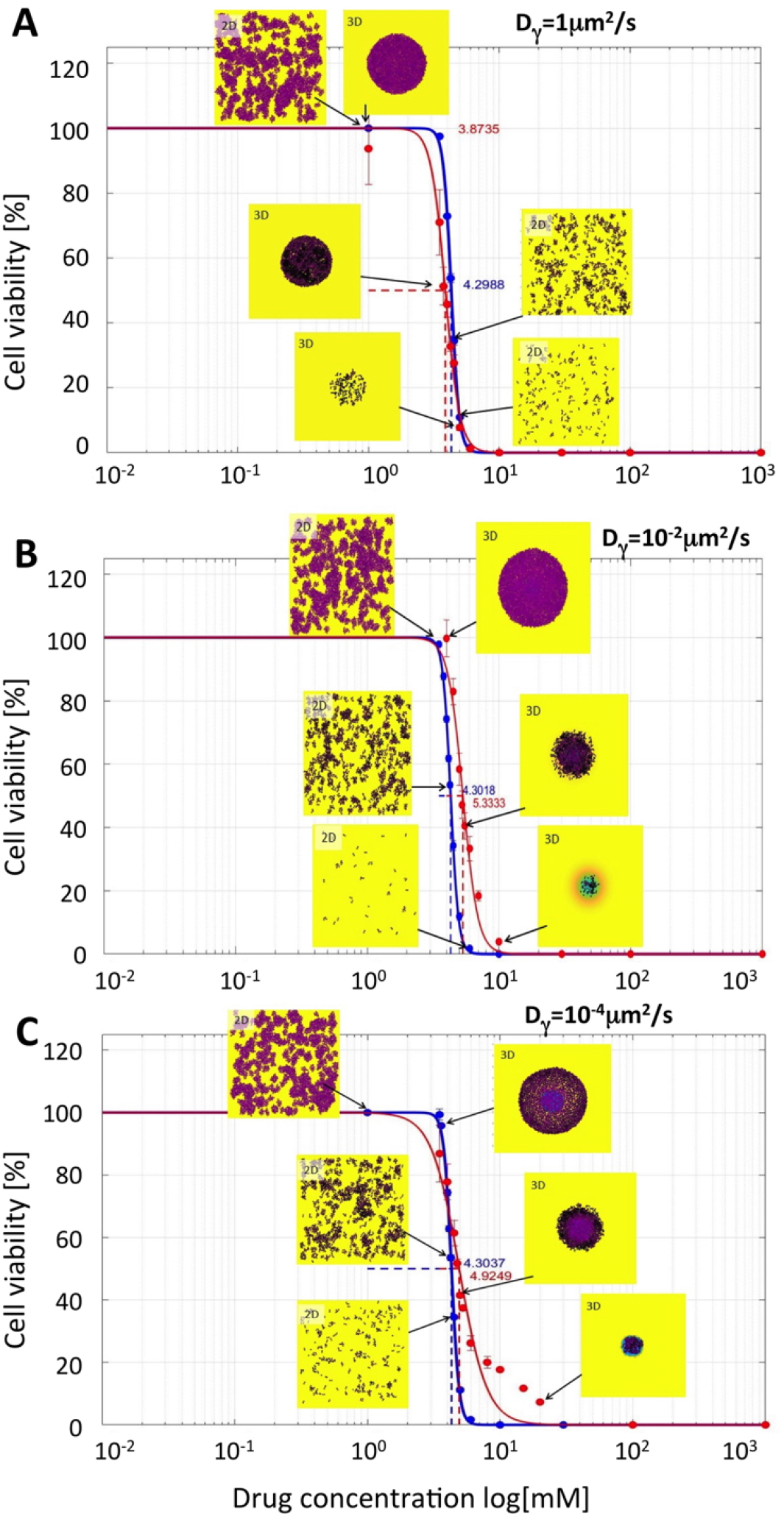
Comparison of IC_50_ curves and values for anti-mitotic drugs. IC_50_ curves for monolayer cultures (blue curves) and cross-sections though spheroid cultures (red curves) and their corresponding IC_50_ values are shown for the following diffusivity values: **A.** D_*γ*_ = 1*μm*^2^/*s*; IC_50_=4.2988 for 2D and 3.8735 for 3D cultures; **B.** D_*γ*_ = 10^−2^*μm*^2^/*s*; IC_50_=4.3018 for 2D and 5.3333 for 3D cultures; **C.** D*_γ_* = 10^−4^*μm*^2^/*s*; IC_50_=4.3037 for 2D and 4.8457 for 3D cultures. The vertical lines show standard deviation values. Insets show final cell configurations for the selected drug concentrations. Cell colors indicate the level of absorbed drug (low-pink, high-black). Background colors indicate the level of the remaining drug (high-yellow, mediumgreen, low-blue). The drug was supplied uniformly only once, at the beginning of each simulation. The cell maturation time for these simulations was 18 hours.

Similarly, as in the cytotoxic drug case, the IC_50_ values for the 2D cell cultures do not differ significantly despite distinct drug diffusivity. Moreover, for fast proliferating cells, the IC_50_ curves for both 2D and 3D cultures are overlapping (Fig. 6A), while for the slowly proliferating cells the IC_50_ curves for the spheroid cultures attain higher half-inhibitory values (Fig. 6BC). In contrast to the cytotoxic case, the 3D cell colonies of cells exposed to higher concentrations of anti-mitotic drugs are less compact. This is an effect of the delayed death of the cells located in the middle of the spheroid. These cells have already absorbed a lethal level of the drug, but remain in the dormant state due to overcrowding. Once the space for their division becomes available, these cells make an attempt to divide and die due to the drug action. This, in turn, creates space for division of other cells, that might instead die if they accumulated a lethal dose of the drug. This may create a domino effect that results in more cell death and less compact spheroid structure. This effect may also take place in 2D cell cultures, but, because these cells are seeded more sparsely, it is not so apparent as in the 3D spherical cultures.

#### The impact of cell maturation ages on IC_50_ values in 2D and 3D cell cultures

Next, we focused on the relationship between the IC_50_ values for the anti-mitotic drugs and cell maturation age. We considered again three cell lines characterized by distinct maturation ages: *A^mat^* = 18, 30 or 50 hours, respectively. These results are summarized in Fig.7A-C where the IC_50_ curves for 2D monolayer culture are shown in red and for the 3D spheroid culture in blue. The corresponding IC_50_ values are summarized in Table 3.

**Figure 7:**
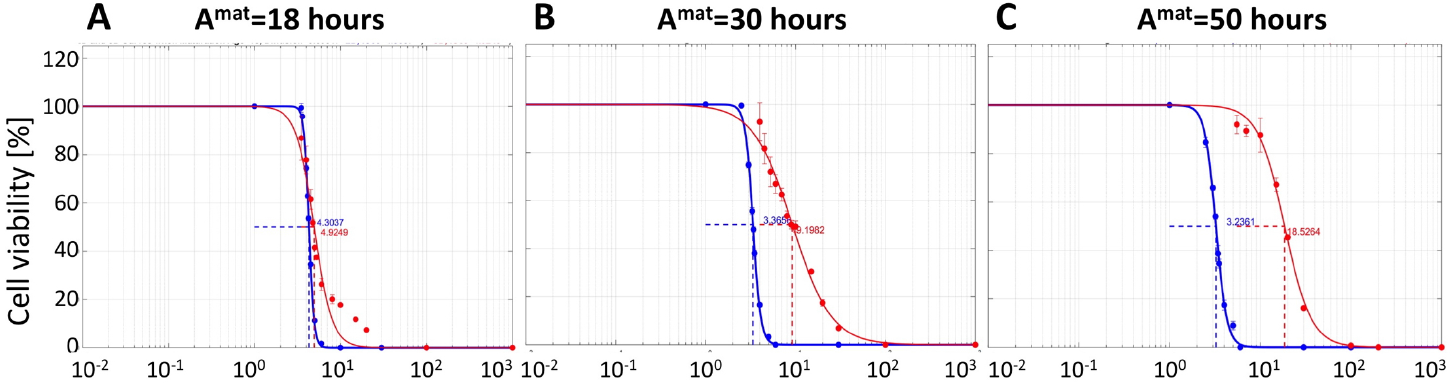
Comparison of IC_50_ curves for three cell lines of different maturation ages. IC_50_ curves for drug diffusion coefficient of *D_γ_* = 10^−4^*μm*^2^/*s* and cell maturation ages of **A.** *A^mat^*=18 hours, **B.** *A^mat^*=30 hours, and **C.** *A^mat^* =50 hours, in 2D cell monolayer culture (blue) and 3D cell spheroid culture (red). All data presented after 72 hours of the simulated time.

**Table 3:** IC_50_ values for anti-mitotic drug diffusion coefficients *D_γ_* and cell maturation ages *A^mat^*, for both monolayer and spheroid *in silico* cultures.

| Summary of $IC_{50}$ values in the 2D Model | | | |
| --- | --- | --- | --- |
| $D_\gamma \mu m^2/s$ | $A^{mat} = 18$ hours | $A^{mat} = 30$ hours | $A^{mat} = 50$ hours |
| 1 | 4.2988 | 3.3495 | 3.2190 |
| $10^{-2}$ | 4.3018 | 3.3641 | 3.1989 |
| $10^{-4}$ | 4.3037 | 3.3656 | 3.2361 |
| Summary of $IC_{50}$ values in the 3D Model | | | |
| $D_\gamma \mu m^2/s$ | $A^{mat} = 18$ hours | $A^{mat} = 30$ hours | $A^{mat} = 50$ hours |
| 1 | 3.8735 | 3.3896 | 3.4942 |
| $10^{-2}$ | 5.3333 | 5.7667 | 7.4052 |
| $10^{-4}$ | 4.8457 | 9.0081 | 18.148 |

We again observe the trend that IC_50_ values increase when drug diffusivity decreases and cell maturation age increases for the 3D cell cultures, however these differences between IC_50_ values and curves are even more pronounced for anti-mitotic drugs in comparison to cytotoxic drugs. In fact, for very slowly proliferating cells (*A^mat^* = 50 hours) there is an order of magnitude difference between IC_50_ values for fast diffusing (*D_γ_* = 1*μm*^2^/*s*) and slowly diffusing (*D_γ_* = 10^−4^*μm*^2^/*s*) anti-mitotic drugs (red curves in Fig. 7A,C). This IC_50_ value is also an order of magnitude larger than the corresponding value for the cytotoxic drugs. In case of the 2D cell monolayer cultures, there is no significant difference in the IC_50_ values despite of differences in drug diffusivity and cell proliferation speed. This is similar to the case of the cytotoxic drugs. For the anti-mitotic drug, we observe a non-monotone relationship between drug diffusivity and IC_50_ values when the average maturation age of the cells is 18 hours (see Table 3, when *A^mat^* = 18 hours). To confirm this non-monotone relationship, we repeated the experiment for the anti-mitotic drug with drug diffusivity *D_γ_* set to 1, 10^−2^ and 10^−4^ *μm*^2^/*s* when *A^mat^* = 18 hours. Results, not shown here, confirm our initial observations.

In summary, the 2D monolayer model is again not able to distinguish between the inhibitory effects of anti-mitotic drugs of different diffusivity for a range of tumor cells with diverse proliferation dynamics. Again, this is attributed to the fact that all cells in the 2D monolayer culture are equally exposed to the drug. In contrast, in the 3D spheroid culture model, both drug diffusivity and cell proliferation time have noticeable effect on the IC_50_ values which increase for slowly diffusing drugs and for slowly proliferating cells. While this trend is similar for cytotoxic and anti-mitotic cells, the overall increase in IC_50_ values is larger for the anti-mitotic cells. Actually, it is an order of magnitude larger for the extreme case (A^*mat*^ = 50 hours and *D_γ_* = 10^−4^*μm*^2^/*s*), showing the crucial role that the drug mechanism of cell killing plays in determining the drug inhibitory effect.

## 6 Discussion

In this paper, we addressed an issue of comparing the potency of anti-cancer drugs using 2D monolayer and 3D spheroid cell cultures. Traditionally, the monolayer cultures are used for evaluation of how a given drug effects growth dynamics of a given cell line or malignant cells derived from a patient’s tumor. These experiments provide information about whether the drug exerts the expected effect and what is the minimal drug concentration needed to observe this effect. The drugs can be then compared to one another using the IC_50_ values which indicate drug concentration that inhibits growth of tumor cell colony by half. By comparing IC_50_ values for different drugs, one can assess which of them is effective at lower concentrations.

However, it has been shown experimentally that the 2D cell cultures do not recreate tumor features observed *in vivo*, such as tumor morphology, cell phenotypes and cell-cell interactions, tumor heterogeneity, and the composition of tumor microenvironment [7, 38, 39]. Therefore, tumor cells’ response to anti-cancer drugs and drug penetration through the tumor tissue might not be faithfully captured in the monolayer cultures. To test this hypothesis, we performed computational studies to compare the IC_50_ curves generated from the analogues of the two-dimensional cell monolayer culture and the three-dimensional multicellular spheroid culture, when the same cells are exposed to the same drugs for the same period of time. We considered hypothetical drugs of various sizes (and thus different diffusivity) and different cell killing mechanisms (cytotoxic and anti-mitotic). Our results indicated that in simulations of 2D cell cultures the IC_50_ values were similar, indicating the same drug potency despite different drug characteristics and cell properties. However, in the simulations of 3D multicellular spheroid cultures, both classes of drugs showed significant differences in the IC_50_ values for drug if different diffusivity and cells of different cell cycles. This strongly suggest that 2D cell cultures are not able to differentiate the half-inhibitory effects for distinct drugs, and the 3D multicellular spheroid cultures should be rather used to assess effective drug concentrations in laboratory experiments.

While in this manuscript, we consider hypothetical drugs of biologically relevant properties, our ultimate goal for our future work is to apply these analyses to specific drugs and specific cell lines/tumors for which we can provide experimental validation. Certain assumptions of our model can also be refined in the future. For example, some cytotoxic drugs can induce cell necrosis while other result in cell lysis. In the former case, cells will remain in the system, though, they would stop absorbing the drug. In the latter case, dying cells may release toxins that have a direct impact on neighboring cells’ survival. The process of dead cell clearance may also be modeled in a more realistic way by using additional time lag, in contrast to instantaneous cell removal, as it is modeled in the current version. Moreover, we observed in our simulations that anti-mitotic drugs may result in less compact spheroid structure due to sudden death of inner cells. This took place when some inner dying cells created space for their immediate neighbors to divide, but the cells died instead when they entered into mitotic phase of their cell cycle. It is observed, that some *in vivo* tumors, such as melanomas, are less dense and contain more interstitial fluid that would suggest more intratumoral death. However, further studies are needed to confirm if the mechanism suggested by our simulations faithfully explains empirical observations.

In summary, to our knowledge, this work is the first comprehensive study comparing drug efficacy in analogues of the 2D and 3D cell cultures. Our simulations indicate that monolayer cell cultures may provide misleading results, since the produced IC_50_ values were almost identical for several cases for which spheroid cultures resulted in significantly distinct IC_50_ values.Thus our main message from this study, is to advocate for using the 3D cell cultures as a standard for testing drug efficacy, which is not currently practiced by the experimental laboratories and pharmacology industry.

## Acknowledgements

This project was initiated as a part of the Preparation for Industrial Careers in Mathematical Sciences (PIC-Math) Award (to NT) from Mathematical Association of America (MAA) and Society of Industrial and Applied Mathematics (SIAM). Support was provided by the National Science Foundation, NSF Grants DMS 1345499 (to NT as PIC-Math Faculty) and DMS 1515442 (to NT). This work was supported by the National Institutes of Health National Cancer Institute grant NIH/NCI U01-CA202229 (to KAR) and the Shared Resources at the H. Lee Moffitt Cancer Center & Research Institute an NCI designated Comprehensive Cancer Center through the National Institutes of Health grant P30-CA076292.

